# The Impact of Electrophysiological Diversity on Pattern Completion in Lithium Nonresponsive Bipolar Disorder: A Computational Modelling Approach

**DOI:** 10.1101/2023.11.20.567781

**Authors:** Abraham Nunes, Selena Singh, Anouar Khayachi, Shani Stern, Thomas Trappenberg, Martin Alda

## Abstract

Patients with bipolar disorder (BD) demonstrate episodic memory deficits, which may be hippocampal-dependent and may be attenuated in lithium responders. Induced pluripotent stem-cell derived CA3 pyramidal cell-like neurons show significant hyperexcitability in lithium responsive BD patients, while lithium nonresponders show marked variance in hyperexcitability. We hypothesize that this variable excitability will impair episodic memory recall, as assessed by cued retrieval (pattern completion) within a computational model of the hippocampal CA3.

We simulated pattern completion tasks using a computational model of the CA3 with different degrees of pyramidal cell excitability variance. Since pyramidal cell excitability variance naturally leads to a mix of hyperexcitability and hypoexcitability, we also examined what fraction (hyper-vs. hypoexcitable) was predominantly responsible for pattern completion errors in our model.

Pyramidal cell excitability variance impaired pattern completion (linear model *β*=-1.94, SE=0.01, p<0.001). The effect was invariant to the number of patterns stored in the network, as well as general inhibitory tone and pyramidal cell sparsity in the network. Excitability variance, and more specifically hyperexcitability, increased the number of spuriously active neurons, increasing false alarm rates and producing pattern completion deficits. Excessive inhibition also induces pattern completion deficits by limiting the number of correctly active neurons during pattern retrieval.

Excitability variance in CA3 pyramidal cell-like neurons observed in lithium nonresponders may predict pattern completion deficits in these patients. These cognitive deficits may not be fully corrected by medications that minimize excitability. Future studies should test our predictions by examining behavioural correlates of pattern completion in lithium responsive and nonresponsive BD patients.

**Author summary:** Patients with bipolar disorder experience debilitating cognitive impairments whose mechanisms are unknown, and these deficits may be greater in patients who do not respond to the mood stabilizer lithium. Studies using induced pluripotent stem cell (iPSC) derived neurons have suggested that CA3 pyramidal cells in lithium nonresponders may have wide diversity of excitability. Our study examines how this diversity of neuronal excitability would impact the computation of pattern completion in the CA3. In a computational model of the CA3, we found that variance in pyramidal cell excitability reliably impaired pattern completion abilities. Furthermore, we found that both the hyperexcitable and hypoexcitable fractions of cells were each responsible for distinct pattern completion errors, depending on the overall level of network inhibition. These results suggest that lithium nonresponsive patients with bipolar disorder will have worse performance on behavioural tasks that are sensitive to pattern completion, potentially including cued recall paradigms. Our results also suggest that mood stabilizers that simply reduce cellular hyperexcitability may not be sufficient to correct micro-circuit level computations in lithium nonresponsive bipolar disorder. Rather, these patients may require development of mood stabilizers that normalize *the distribution* of neuronal hyperexcitability among CA3 pyramidal cells.

## Introduction

Bipolar disorder (BD) is a chronic and debilitating mental illness characterized by episodes of mania and depression [1], whose neurobiology remains unknown [2]. A key observation in the clinical management of BD is the variability in response to lithium [3], which has been the gold-standard prophylactic mood stabilizer for more than 60 years [4]. While lithium effectively mitigates mood symptoms for a sizable minority of patients, approximately two-thirds of patients remain nonresponsive [3]. Intriguingly, lithium nonresponders may have particularly poor cognitive functioning compared to patients who are stable on lithium monotherapy, especially in the domain of episodic memory [5]. This raises important questions about the underlying neural mechanisms of episodic memory functioning in BD, and how these mechanisms relate to treatment responsiveness.

The hippocampal CA3 region is known for its role in memory processing, particularly in rapid one-shot learning [6] and pattern completion [7]. Pattern completion is the ability to retrieve a complete memory representation from any of that memory’s parts [8]. This has been postulated to occur in the hippocampal CA3 by virtue of the extensive recurrent collateral connections in CA3 [9–14]. The recurrent collateral synapses in CA3 facilitate pattern completion by allowing pyramidal cells representing some sub-component of a memory to activate other neurons representing the same memory, ultimately completing the representation from only partial cues. These operations suggest that the CA3 may perform pattern completion by functioning as an autoassociative attractor network [9, 11]. The hypothesis that CA3 performs pattern completion has empirical support [7, 15, 16], and further modelling studies have gone on to examine network storage capacity and the influence of inhibitory interneurons and network sparsity [12, 14, 17], connectivity patterns [18], and symmetry of plasticity mechanisms [19] on pattern completion, to name a few. Pattern completion may be probed behaviourally by having participants first encode a set of stimuli, and subsequently (A) recall old stimuli or (B) discriminate old from lure stimuli based on partial or degraded/noisy cues [20]. Pattern completion is believed to be disrupted in psychiatric conditions such as schizophrenia, and may have a role in the formation of delusions [21–23]. Given the genetic relatedness of BD and schizophrenia [24], and BD’s association with both psychosis [25] and declarative memory impairments [5, 26–29], it is plausible that similar hypothesized pattern completion deficits may be found in BD. Several lines of research suggest that CA3 structure, function, and electrophysiology may be impaired in BD, which we review below.

First, many genetic variants associated with BD and lithium responsiveness are also implicated in CA3 structure and function. The GRIN1 gene, which encodes the NR1 subunit of the N-methyl-D-aspartate receptor (NMDAR) is associated with and downregulated in BD [30, 31] (but see [32]) and schizophrenia [33]. Animal models demonstrate that GRIN1 mediates the integrity of conjunctive and associative representations in the CA3 [15, 34]. In addition to the BDNF-NTRK2 pathway being associated with BD [31], completed suicide [35], and response to mood stabilizers [36–39] at the behavioural level, its involvement extends down to the cellular and circuit level as well, regulating dendritic spine density in CA3 [40] and the establishment of functional circuitry between the dentate gyrus and CA3 [41]. While these genetic abnormalities may also predispose broader neurobiological changes in BD, their overlap with CA3 structure and functioning collectively suggest that studying this brain region in BD is an important research direction.

In addition to genetic abnormalities, functional impairment may also be attributable in part to reduced hippocampal volume in BD. The largest analysis of hippocampal subfield volumes in BD to date (1472 patients and 3226 controls) has found significantly smaller CA3 volume in BD patients (Cohen’s d=-0.20) [42]: an abnormality which may be associated with impaired memory recall [43]. Interestingly, lithium users showed greater preservation of CA3 volume compared to lithium nonusers (n=464) [42]. Reductions in hippocampal volume may be explained by reductions in parvalbumin-positive interneuron number and size [44, 45], which may impact the regulation of the hyperexcitable fraction of pyramidal cells in the CA3.

Recent advances in stem cell technology have allowed for the precise investigation of the properties of patient-derived *in vitro* models of CA3 pyramidal cells, to further elucidate potential cellular-level abnormalities in BD [46]. Specifically, to study the electrophysiological properties associated with lithium responsiveness, patient-derived cells have been reprogrammed into induced pluripotent stem cells (iPSCs), and subsequently differentiated into CA3 pyramidal cell-like neurons (CA3-PCs) [47]. Notably, CA3-PCs derived from lithium responders are hyperexcitable, which is normalized upon lithium exposure [47]. This phenomenon is absent in CA3-PCs from both healthy controls and lithium nonresponders [47]. Yet, CA3-PCs derived from lithium nonresponders have exhibited high diversity of activity, with a mixed population of hyperexcitable and hypoexcitable cells [47, 48]. This electrophysiological heterogeneity is a distinct abnormality between lithium responders and nonresponders and may be a potential key to understanding the neural underpinnings of cognitive dysfunction in lithium nonresponsive BD.

Together, there is genetic, structural, and cellular electrophysiological evidence suggesting that CA3 structure and functioning are likely to be abnormal in BD. However, to link these abnormalities to observable behaviours, we must understand (A) the computations carried out by the CA3 circuit, (B) how these computations are affected by the neurobiological abnormalities observed in BD, and (C) how these computations connect to observable behaviours. As a first step, we must gain an understanding of how variability in cellular excitability in the CA3 relates to circuit-level computations. Therefore, in this study, we leverage computational modeling to examine how the diversity of excitability in CA3-PCs might affect pattern completion in the CA3. This work will facilitate our ability to bridge the gap between cellular properties and network-level function in BD, providing more specific predictions about the memory dysfunctions seen in lithium nonresponsive BD.

## Results

Our study extends a previously developed model of the hippocampal CA3 suitable for large-scale simulations [12, 14, 18, 19]. We use an implementation that includes *n*=3,000 integrate-and-fire-type glutamatergic neurons equipped with a symmetric plasticity rule (Equation 3) and a pooled inhibitory population modelled as a single unit (Fig 1). Each pyramidal cell had a unique inhibitory scaling factor 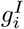, which facilitated the modelling of variable levels of excitability. We assumed a random (Erdős–Rényi) connectivity structure with probability *c*_*ij*_ = *c*^*^, where *c*_*ij*_ is the probability of an excitatory synaptic connection from neuron *i* to neuron *j*. Pattern completion behaviour in this model has been previously shown to be comparable to that in a larger-scale implementation of 330,000 cells [18, 19].

**Fig 1.**
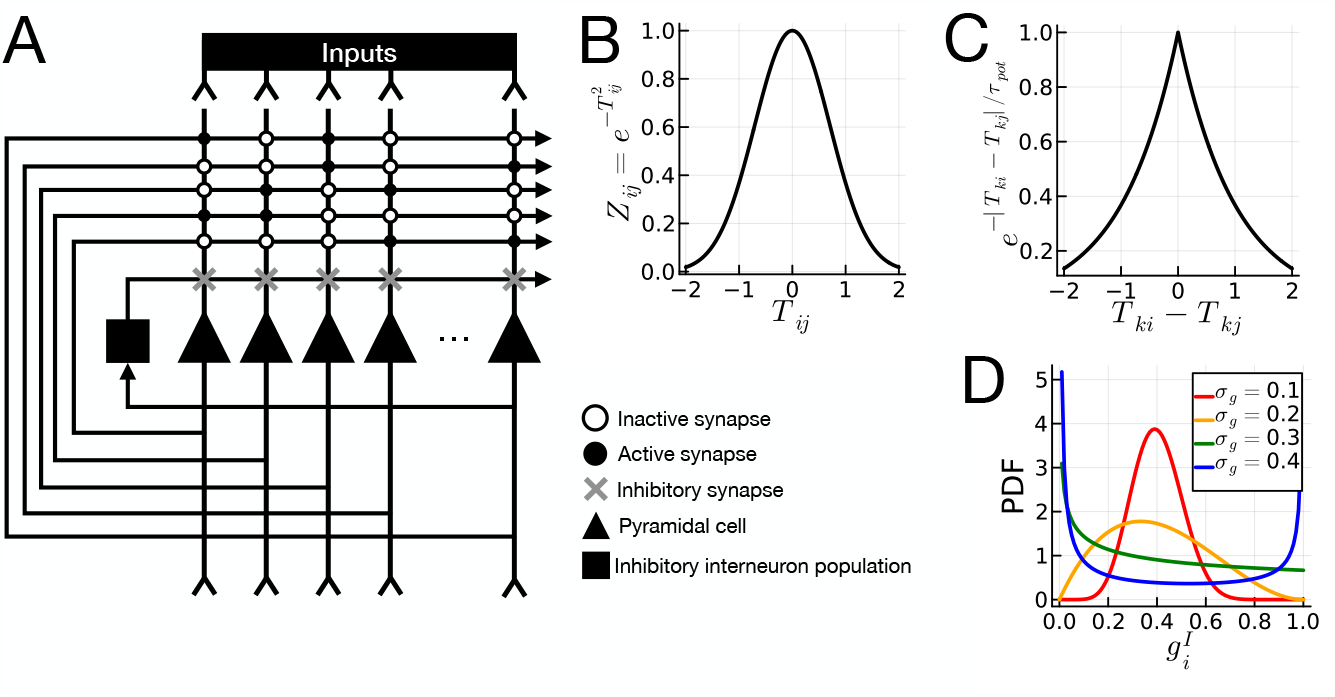
Illustration of the computational model. *Panel A:* Model architecture. Solid triangles represent the CA3 pyramidal cells. Solid square is the inhibitory interneuron population. Solid circles are active synapses at which plasticity occurs. Open circles are inactive synapses, which effectively represent no connection or plasticity between two neurons. Gray “x” markers are inhibitory synapses. Note that while there are inhibitory inputs to all pyramidal cells, they will vary in strength depending on the value of 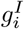. ***Panel B:*** Relationship between spike time *Tij* and activation level *Zij*. ***Panel C:*** Symmetric spike timing-dependent plasticity function [19, 49]. The x-axis plots the difference in spike time between neurons *i* and *j* during pattern *k*, denoted *Tki*-*Tkj*, and the y-axis denotes the resulting degree of synaptic potentiation, exp{-|*Tki*-*Tkj*| */t*pot, which applies only at synapses that are connected (that is, “active synapses”). ***Panel D:*** Illustration of Beta distribution with *µg* = 0.4, and *σg* ∈ {0.1, 0.2, 0.3, 0.4}. The X-axis shows the value of 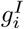, and the y-axis is the probability density.

### Hyperexcitability and Diverse Excitability Impair Pattern Completion

We are interested in examining pattern completion performance in the CA3 under conditions of different degrees and variability of pyramidal cell excitability. During the encoding phase, a set of *m* ∈ N_+_ patterns, each characterized by temporally distributed activity in a proportion 0 *< a <* 1 of neurons, were applied to the network, which engaged in learning via symmetric spike-timing dependent plasticity (Equation 3; Fig 1). During the retrieval phase, a partial “seed” pattern was presented to the CA3 network as a cue for recall. Each seed pattern corresponded to one of the *m* patterns but with only a proportion 0 *< b*_1_ *<* 1 of the original activations intact. The network was then allowed to equilibrate over 10 iterations to recover the input pattern.

We measured pattern completion performance as the Pearson correlation *ρ*(*X*_*i*_, *Z*_*i*_) between the recovered pattern *X*_*i*_ = (*X*_*ij*_)_*j*=1,2,…,*n*_ and its corresponding ground truth pattern *Z*_*i*_ = (*Z*_*ij*_)_*j*=1,2,…,*n*_. Our independent variable of interest is the standard deviation of the cell-specific inhibitory scaling factor 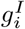, denoted *σ*_*g*_. A wider distribution on 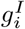 simulates the heterogeneous excitability observed in lithium nonresponsive BD [47, 48]. The values 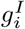 are sampled from a Beta distribution with mean *µ*_*g*_ and standard deviation *σ*_*g*_, where 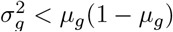. We examine the pattern completion ability of the network while systematically varying *σ*_*g*_, conditional upon *m, µ*_*g*_, *a*, and *c*^*^ for neurons (*i, j*) = 1, 2, …, *n* . We measured the effect of *σ*_*g*_ on pattern completion performance using linear regression of *ρ* against *σ*_*g*_, with *m, µ*_*g*_, *a*, and *c*^*^ as covariates.

Table 1 and Fig 2 show the effects of hyperexcitability, and diversity thereof, on pattern completion performance in our CA3 model. Diverse pyramidal cell excitability, measured by *σ*_*g*_, reduced pattern completion performance in all cases (top row of plots in Fig 2; *β*=-1.94, SE=0.01, p*<*0.001, Table 1). The effect was invariant to pattern load (*m*), evinced by the relatively parallel decline in correlation across values of *m* in Fig 2. Hyperexcitability, captured by higher values of *a* (the proportion of neurons active in a given pattern; *β*=-2.89, SE=0.01, p*<*0.001), as well as lower values of *µ*_*g*_ (*β*=0.74, SE=0.01, p*<*0.001), was associated with worse pattern completion performance. The impact of variability of excitability *σ*_*g*_ was independent of overall levels of inhibition *µ*_*g*_ or the proportion of neurons active in each pattern (*a*), suggesting that correction of overall levels of excitability is insufficient to fully correct pattern completion deficits in the presence of variable excitability of CA3 pyramidal cells (S1 Fig, S2 Fig, S3 Fig).

**Table 1.** Ordinary least squares estimates of effects of diversity of hyperexcitability (*σ*_*g*_), as well as overall levels of excitability (the proportion of neurons active for each pattern, *a*, and the average inhibitory strength across neurons, *µ*_*g*_), controlled for the overall pattern load (*m*) and connectivity rate (*c*^*^). Abbreviation: regression coefficient (*β*), standard error (SE), t-statistic (t), p-value (p), confidence interval (CI).

| | $\beta$ | SE | t | p | 95% CI Low | 95% CI High |
| --- | --- | --- | --- | --- | --- | --- |
| Intercept | 0.65 | 0.004 | 180.09 | < 0.001 | 0.64 | 0.65 |
| $m$ | -0.004 | 4.52e<sup>-5</sup> | -90.77 | < 0.001 | -0.004 | -0.004 |
| $a$ | -2.89 | 0.01 | -285.61 | < 0.001 | -2.91 | -2.87 |
| $c^*$ | 0.17 | 0.003 | 47.15 | < 0.001 | 0.16 | 0.17 |
| $\mu_g$ | 0.74 | 0.01 | 101.70 | < 0.001 | 0.73 | 0.76 |
| $\sigma_g$ | -1.94 | 0.01 | -201.82 | < 0.001 | -1.96 | -1.92 |

**Fig 2.**
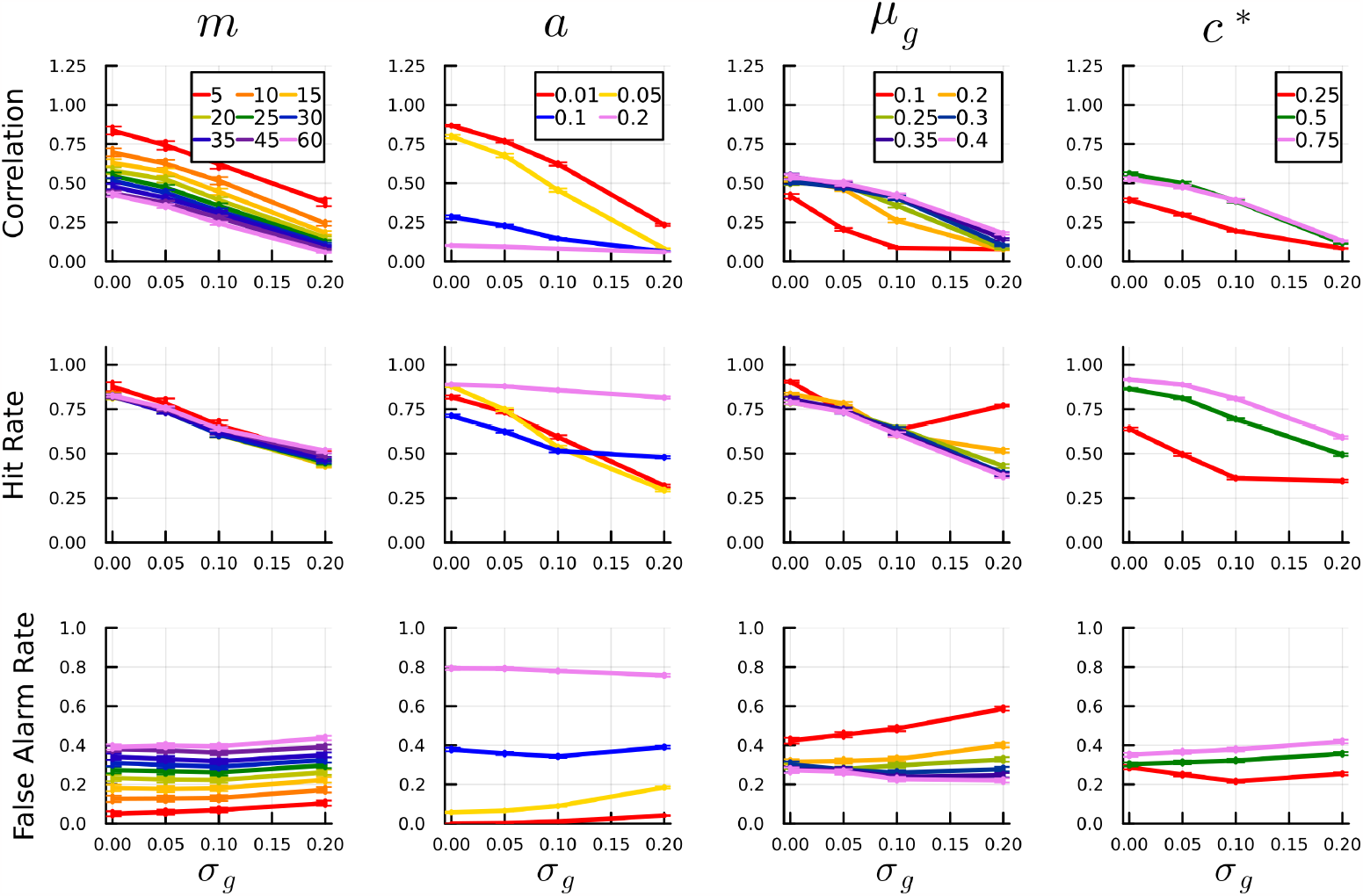
Effects of excitability variability on pattern completion performance. Pattern completion performance is measured by the Pearson correlation between the true and recovered patterns. The top row of plots shows correlation results, while the second and third rows show the hit rates and false alarm rates with respect to *σ*_*g*_. Columns display results with respect to various moderating factors, including the number of encoded patterns (*m*), the proportion of neurons active for each pattern (*a*), the mean level of inhibition (*µ*_*g*_), and average connectivity (*c*^*^).

As secondary outcomes, we examined the amount of valid and spurious activity during recall. Valid activity is computed as the hit rate

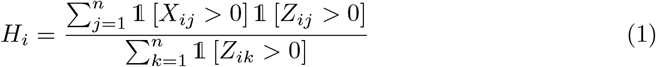

where 𝟙[*x*] is an indicator function taking values 1 if *x* is true, and 0 otherwise. Spurious activity was computed as the false alarm rate

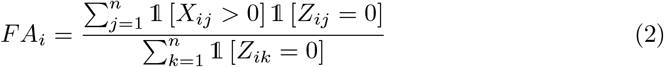

which is the propensity for a given pyramidal cell neuron in pattern *i* to be active during recall, despite not being involved in the encoded pattern *Z*_*i*_.

Hyperexcitability impaired pattern completion performance largely by increasing false alarm rates (that is, increasing the number of spuriously active neurons), while variability of excitability (*σ*_*g*_) impaired pattern completion largely by reducing hit rates (Fig 2).

### Variable Excitability Induces Pattern Completion Errors Through the Hyperexcitable Cells

We then probed the degree to which pattern completion errors were attributable to the hyperexcitable vs. hypoexcitable fraction of pyramidal cells. Across multiple average levels of inhibition (*µ*_*g*_∈ {0.1, 0.2, 0.3, 0.4, 0.5, 0.6}), we examined the pattern completion error rate in relation to pyramidal cell inhibition variance 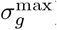 in increments of 0.02, with 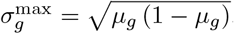.

Fig 3 shows the results of an evaluation of error rates in pattern completion in relation to the variability of excitability levels (*σ*_*g*_) across multiple mean inhibition levels (*µ*_*g*_). Across all mean inhibition levels, increases in variability of excitability resulted in higher pattern completion error rates primarily attributable to the hyperexcitable half of the CA3 pyramidal cell population. At low levels of inhibition (*µ*_*g*_=0.1; high global levels of hyperexcitability), the lower bound on error rates is approximately 0.25, The lower bound on error rate subsequently decreases to almost 0 before increasing to an upper bound of 0.10 for high levels of inhibition (*µ*_*g*_ above 0.4). At high levels of inhibition, the pattern completion error rate was constant for the hypoexcitable fraction of pyramidal cells.

**Fig 3.**
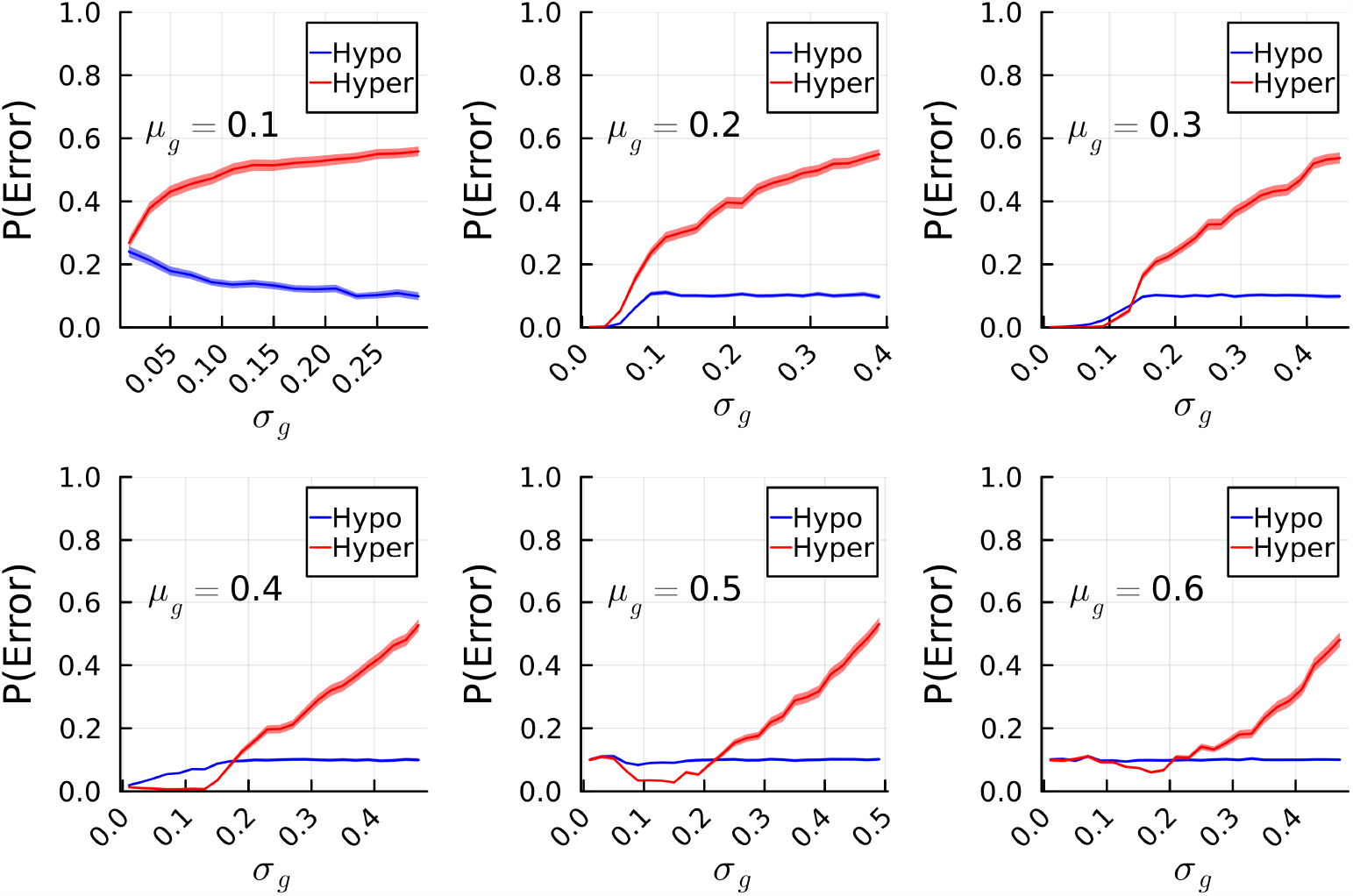
Relationship of error rates to hyper/hypo-excitable fractions. Pattern completion error rates (y-axes) in hyperexcitable (red lines; inhibition level less than mean *µ*_*g*_) and hypoexcitable (blue lines; inhibition level less higher than mean *µ*_*g*_) neurons, in relation to the level of variability in pyramidal cell activity (x-axes; *σ*_*g*_). Each plot corresponds to a specific mean level of inhibition (*µ*_*g*_). Solid lines are mean error rates, and ribbons are 95% confidence intervals.

## Discussion

The present study demonstrates that variability of CA3 pyramidal cell excitability, which has been observed in iPSC-derived CA3 pyramidal cell-like neurons from lithium nonresponders, impairs pattern completion in an autoassociative attractor network model of the CA3. The impairment of pattern completion secondary to variable excitability was independent of average population-level excitability, and was invariant to any other manipulation of network architecture or dynamics, suggesting that variability in iPSC-derived pyramidal cell activity is potentially a candidate independent marker of episodic memory impairments in lithium nonresponders. Pattern completion errors arising from variability of neuronal excitability arose mainly because some cells were more hyperexcitable than others. However, under some circumstances (that is, when the overall average excitability levels are low), pattern completion errors arise primarily from neurons that are hypoexcitable relative to the rest of the population. Our findings predict that mechanisms of cognitive impairment related to pattern completion and episodic memory in lithium nonresponders may have physiologically distinct mechanisms compared to impairments arising in lithium responders.

Furthermore, our results suggest that remedying pattern completion deficits in lithium responders and nonresponders, should they be identified experimentally, may require different approaches.

The 50% most excitable neurons (that is, the hyperexcitable fraction) substantially contributed to pattern completion errors. However, when overall excitability levels were low (high inhibition, corresponding to high *μ*), we observed an increase in the minimum error rate driven primarily by the 50% least excitable neurons (that is, the hypoexcitable fraction). This increase in error rates in response to increasing inhibition predicts that lithium nonresponsive patients receiving excitability-lowering mood stabilizers, potentially including anticonvulsants [46, 50], may exhibit a ceiling effect on their cognitive performance with treatment. In other words, reduction of pyramidal cell hyperexcitability may improve memory performance to a point, after which it may worsen again. These results highlight the importance of understanding how mood stabilizers may normalize the distribution of neuronal excitability, rather than merely reducing hyperexcitability overall. The complexity of this type of intervention stands in contrast to the potential effects of treatment on lithium responders, whose CA3 pyramidal cells do not show a wide variation in hyperexcitability levels [47, 48]. For lithium responders, simply limiting cellular hyperexcitability would be predicted to improve pattern completion. Future studies should examine behavioural measures of pattern completion in lithium responders and nonresponders from whom iPSC-derived CA3 pyramidal cell-like neurons have been cultured, and whose excitability distributions (mean excitability and variance) are well characterized. Such studies should then characterize how neuronal excitability distributions are affected when the neuronal cultures are exposed to mood stabilizers to which patients demonstrably respond or fail to respond. Our results predict that persistent variance in excitability despite treatment with excitability-lowering mood stabilizers will be associated with worse behavioural pattern completion performance in patients with BD.

Our results also emphasize the importance of understanding the mechanisms underlying the diversity of neuronal excitability in lithium nonresponders. Stern et al. [47, 48, 51] showed that BD nonresponders have reduced sodium currents on average as compared to the neurons derived from healthy individuals. This reduction in sodium currents is paired with an increase of several types of potassium currents. Differences in sodium currents were found to potentially mediate variation between hyperexcitable and hypoexcitable neurons from lithium nonresponders; the neurons with the average or high sodium currents were mostly hyperexcitable, while the neurons with sodium currents below the average (which is already lower in the nonresponders compared to both controls and the responders) were mostly hypoexcitable and unable to produce action potentials even in response to current injections. We believe that this reduction in the sodium currents that is paired with an increase in potassium currents may have a strong influence on the BD nonresponders’ neuronal electrophysiological diversity. However, the underlying cause of this diversity in sodium and potassium conductances within individual patients is unknown. Upon application of chronic lithium treatment, neurons derived from the BD lithium responders, increased their sodium currents and decreased their potassium currents, “normalizing” their neurophysiology and making them more like control neurons [47, 51]. Lithium even had the effect of normalizing neuronal morphology. In contrast, when treating the neurons with valproic acid [50], both neurons derived from BD responders and the nonresponders, exhibited a reduction in their sodium currents. This drives neurons derived from BD nonresponders even further away from the normal neurophysiology [52], further supporting the notion that using medication to simply decrease hyperexcitability may not be the optimal treatment and the overall neurophysiological features of the neurons and the network that they form should be considered.

A major strength of our study is the simplified and well-controlled nature of our computational model of the CA3. This facilitated precise control over the overall and mean levels of excitability variance in the CA3 pyramidal cell population. The simplicity of this model facilitated the testing of many potential confounding factors, ultimately demonstrating that the relationship between variance in CA3 pyramidal cell excitability and pattern completion deficits were invariant to overall levels of inhibition, connectivity levels, pattern load, and network sparsity. However, the simplicity of this model is also a limitation, given that it is (by definition) an abstraction of real-world neuronal networks. For example, our model does not include the large diversity of interneuron types in the CA3 that may be considered in more detailed biophysical models [22]. The leaky integrate and fire point neurons used in our model also do not capture the nuances of the structural and biophysical properties of the somatodendritic tree of CA3 pyramidal cells [40, 41]. Although we could not identify a specific level of experimentally-determined variance in CA3 pyramidal cell-like excitability, our results are robust to this because we examined pattern completion performance across the full range of excitability variance available under our model. Our model also employs a relatively simple and dense connectivity pattern between CA3 pyramidal cells. The connectivity patterns between CA3 pyramidal cells have previously been shown to incorporate complex motifs [18], although simpler dense and random connectivity patterns as employed in the present study have been shown to generate similar network behaviour [19]. To efficiently use computational resources, we employed the simpler approach, which facilitated our examination of different network conditions in sensitivity analyses.

## Conclusion

In conclusion, our study suggests that the diverse excitability of CA3 pyramidal cell-like neurons observed in lithium nonresponders may predict pattern completion deficits in these patients. These pattern completion deficits are invariant to overall excitability levels, suggesting that even if overall CA3 pyramidal cell excitability is controlled in lithium nonresponsive BD patients, these patients may continue to exhibit cognitive deficits that depend on pattern completion. These predictions should be validated experimentally using behavioural paradigms that require patients to first encode a set of stimuli, after which they must either (A) recall those stimuli or (B) discriminate the studied stimuli from novel/lure stimuli, given partial or noisy/corrupted cues. Such paradigms exist [20], and should be used to study behavioural pattern completion in lithium responsive and nonresponsive patients, in parallel with excitability variance investigations using iPSC-derived CA3 pyramidal cell-like neurons derived from the same patients. Furthermore, it would be useful to examine these behavioural pattern completion deficits and how they correlate to both clinical and cellular responses to lithium or anticonvulsant mood stabilizers. Our results suggest that without normalizing the complete distribution of cellular excitability in the CA3, pattern completion deficits may persist in patients with BD.

## Materials and Methods

### Computational Model

#### Encoding Phase and Learning in the Model

For a given storage cycle, a pattern *i* ∈ {1, 2, …, *m*} is presented to the CA3. Each pattern is characterized by activity in *n*_*a*_ = *n* × *a*⌋ neurons, where *a* ∈ ⌊0, 1] is the proportion of neurons active in a given pattern, and ⌊*x*⌋ is the floor function.

For each pattern *i* ∈*{*1, 2, …, *m}* presented to the CA3 in a given storage cycle, let *T*_*ij*_ ∼Normal(0, *s*_*T*_) be the spiking time of neuron *j* ∈*{*1, 2, …, *n}*, with mean 0, and standard deviation *s*_*T*_ *>* 0. Following Mishra et al. [19] we set *s*_*T*_ = 0.2, corresponding to 20% of a storage cycle. The activation level of neuron *j* in pattern *i* is 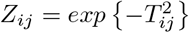 (Fig 1). For the excitatory synapse from neuron *j* to *i*, plasticity during the encoding phase was implemented using the following symmetric spike timing-dependent plasticity rule [19]:

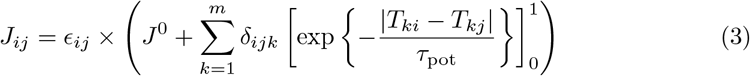

where *ϵ*_*ij*_ ∼Bernoulli(*c*_*ij*_) is a binary random variable with probability *c*_*ij*_ indicating whether there is an excitatory connection from neuron *j* to *i, δ*_*ijk*_ is an indicator variable denoting whether both neurons *i* and *j* are active during pattern *k*, and 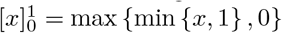 . The value *J*^0^ = 0 indicates that the base synaptic strength is 0. The synaptic plasticity time constant *τ*_pot_ was set to 1, corresponding to a single storage cycle [19].

### Pattern Completion Task and Recall Dynamics in the Model

After a CA3 network learns patterns 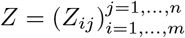, we evaluated pattern completion by presenting the network with an incomplete/corrupted version of each pattern in *Z*, and evaluating the accuracy with which the CA3 network could recover the original pattern.

Dynamics of pattern completion during recall were simulated by feeding a partial or noisy “seed” input pattern to the CA3 network, and allowing network dynamics to equilibrate over 10 recall cycles. The seed pattern corresponding to pattern *k* = {1, 2, …, *m}* is controlled by parameters, *b*_1_ ∈ [0, 1] (the proportion of neurons in the seed pattern that correspond to valid activations in pattern *k*) and *b*_2_ ∈ [0, 1] (the proportion of spurious additional activations that were not present in the true pattern *k*). Increasing *b*_1_ corresponds to providing the network with a seed pattern that is more accurately reflective of a true pattern upon which the synapses were trained during encoding. Increasing *b*_2_ corresponds to adding more “noise” in the form of spuriously active neurons. In the present study, as in Mishra et al., [19] we set *b*_1_ = 0.5, and *b*_2_ = 0.

The total synaptic input into neuron *i* at recall cycle *t* is:

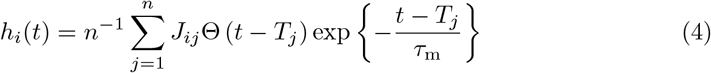

where Θ(*x*) is the Heaviside step function, which takes a value of 1 for *x >* 0, and 0 otherwise, *τ*_m_ is the time constant for neuron *i*’s membrane potential (*τ*_m_ is assumed to be the same for all pyramidal cells in the model), and *T*_*j*_ is the spike time of neuron *j* at the previous recall cycle. Neuron *i* spikes as part of pattern *k* 1, 2, …, *m* if 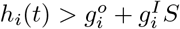, where *S* is the sum of all neural activity at the previous recall cycle, 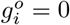 is the basal action potential threshold for neuron *i*, and 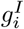 is a factor scaling the degree of inhibition received by neuron *i. S* models a general inhibitory population whose activity is proportional to the total network activity at the previous recall cycle. All simulations were conducted using custom scripts in the Julia programming language (v. 1.9.0), which are available as supplementary material (S1 File).

## Supporting information

**S1 Fig. Interactions between model parameters and pattern correlation**. Activity correlation (y-axis) and interactions between the diversity of hyperexcitability (*σ*_*g*_; x-axis), activation level (a), and mean level of inhibition (*μ*_*g*_). Coloured lines represent the pattern load, ranging from *m*=5 (red) to *m*=65 (indigo), in steps of 5, according to a rainbow palette.

**S2 Fig. Interactions between model parameters and hit rates**. Hit rates (y-axis) and interactions between the diversity of hyperexcitability (*σ*_*g*_; x-axis), activation level (*a*), and mean level of inhibition (*μ*_*g*_). Coloured lines represent the pattern load, ranging from *m*=5 (red) to *m*=65 (indigo), in steps of 5, according to a rainbow palette.

**S3 Fig. Interactions between model parameters and false alarm rates**. False alarm rates (y-axis) and interactions between the diversity of hyperexcitability (*σ*_*g*_; axis), activation level (*a*), and mean level of inhibition (*μ*_*g*_). Coloured lines represent the pattern load, ranging from *m*=5 (red) to *m*=65 (indigo), in steps of 5, according to a rainbow palette.

**S1 File. Code repository to reproduce experiments and analyses**.

## References

1. Grande I, Berk M, Birmaher B, Vieta E. Bipolar disorder. The Lancet. 2015;6736(15):1–12. doi:10.1016/S0140-6736(15)00241-X.

2. American Psychiatric Association. Diagnostic and Statistical Manual of Mental Disorders, 5th Edition (DSM-5). American Psychiatric Publishing; 2013.

3. Garnham J, Munro A, Slaney C, MacDougall M, Passmore M, Duffy A, et al. Prophylactic treatment response in bipolar disorder: Results of a naturalistic observation study.Journal of Affective Disorders. 2007;104(1):185–190. doi:10.1016/j.jad.2007.03.003.

4. Grof P. Sixty years of lithium responders. Neuropsychobiology. 2010;62(1):8–16.

5. Burdick KE, Millett CE, Russo M, Alda M, Alliey-Rodriguez N, Anand A, et al. The association between lithium use and neurocognitive performance in patients with bipolar disorder. Neuropsychopharmacology. 2020;45(10):1743–1749. doi:10.1038/s41386-020-0683-2.

6. Nakashiba T, Young JZ, McHugh TJ, Buhl DL, Tonegawa S. Transgenic Inhibition of Synaptic Transmission Reveals Role of CA3 Output in Hippocampal Learning. Science. 2008;319(5867):1260–1264. doi:10.1126/science.1151120.

7. Neunuebel JP, Knierim JJ. CA3 Retrieves Coherent Representations from Degraded Input: Direct Evidence for CA3 Pattern Completion and Dentate Gyrus Pattern Separation. Neuron. 2014;81(2):416–427. doi:10.1016/j.neuron.2013.11.017.

8. Rolls ET. The mechanisms for pattern completion and pattern separation in the hippocampus. Frontiers in Systems Neuroscience. 2013;7:74. doi:10.3389/fnsys.2013.00074.

9. Marr D. Simple memory: a theory for archicortex. Philosophical Transactions of the Royal Society of London B, Biological Sciences. 1971;262(841):23–81. doi:10.1098/rstb.1971.0078.

10. Rolls ET. Information representation, processing and storage in the brain: Analysis at the single neuron level. In: Changeux JP, Konishi M, editors. The neural and molecular bases of learning.Chichester: Wiley; 1987. p. 503–540. Available from: https://scholar.google.com/scholar_lookup?title=Information%20representation%2C%20processing%20and%20storage%20in%20the%20brain%3A%20Analysis%20at%20the%20single%20neuron%20level&publication_year=1987&author=E.T.%20Rolls.

11. McNaughton BL, Morris RGM. Hippocampal synaptic enhancement and information storage within a distributed memory system. Trends in Neurosciences. 1987;10(10):408–415. doi:10.1016/0166-2236(87)90011-7.

12. Gibson WG, Robinson J. Statistical analysis of the dynamics of a sparse associative memory. Neural Networks. 1992;5(4):645–661. doi:10.1016/S0893-6080(05)80042-5.

13. Rolls ET. Pattern separation, completion, and categorisation in the hippocampus and neocortex. Neurobiology of Learning and Memory. 2016;129:4–28. doi:10.1016/j.nlm.2015.07.008.

14. Bennett MR, Gibson WG, Robinson J. Dynamics of the CA3 Pyramidal Neuron Autoassociative Memory Network in the Hippocampus. 2021; p. 22.

15. Nakazawa K, Quirk MC, Chitwood RA, Watanabe M, Yeckel MF, Sun LD, et al. Requirement for Hippocampal CA3 NMDA Receptors in Associative Memory Recall. Science. 2002;297(5579):211–218. doi:10.1126/science.1071795.

16. Gold AE, Kesner RP. The role of the CA3 subregion of the dorsal hippocampus in spatial pattern completion in the rat. Hippocampus. 2005;15(6):808–814. doi:10.1002/hipo.20103.

17. Treves A, Rolls ET. What determines the capacity of autoassociative memories in the brain? Network: Computation in Neural Systems. 1991;2(4):371–397. doi:10.1088/0954-898X24004.

18. Guzman SJ, Schlögl A, Frotscher M, Jonas P. Synaptic mechanisms of pattern completion in the hippocampal CA3 network. Science. 2016;353(6304):1117–1123. doi:10.1126/science.aaf1836.

19. Mishra RK, Kim S, Guzman SJ, Jonas P. Symmetric spike timing-dependent plasticity at CA3–CA3 synapses optimizes storage and recall in autoassociative networks. Nature Communications. 2016;7(1):11552. doi:10.1038/ncomms11552.

20. Liu KY, Gould RL, Coulson MC, Ward EV, Howard RJ. Tests of pattern separation and pattern completion in humans—A systematic review. Hippocampus. 2016;26(6):705–717. doi:10.1002/hipo.22561.

21. Tamminga CA, Stan AD, Wagner AD. The Hippocampal Formation in Schizophrenia. American Journal of Psychiatry. 2010;167(10):1178–1193. doi:10.1176/appi.ajp.2010.09081187.

22. Neymotin SA, Lazarewicz MT, Sherif M, Contreras D, Finkel LH, Lytton WW. Ketamine Disrupts Theta Modulation of Gamma in a Computer Model of Hippocampus. Journal of Neuroscience. 2011;31(32):11733–11743. doi:10.1523/JNEUROSCI.0501-11.2011.

23. Tamminga CA, Southcott S, Sacco C, Wagner AD, Ghose S. Glutamate Dysfunction in Hippocampus: Relevance of Dentate Gyrus and CA3 Signaling. Schizophrenia Bulletin. 2012;38(5):927–935. doi:10.1093/schbul/sbs062.

24. Anttila V, Bulik-Sullivan B, Finucane HK, Walters RK, Bras J, Duncan L, et al. Analysis of shared heritability in common disorders of the brain. Science. 2018;360(6395):eaap8757. doi:10.1126/science.aap8757.

25. Goes FS, Sadler B, Toolan J, Zamoiski RD, Mondimore FM, MacKinnon DF, et al. Psychotic features in bipolar and unipolar depression. Bipolar Disorders. 2007;9(8):901–906. doi:10.1111/j.1399-5618.2007.00460.x.

26. Cardenas SA, Kassem L, Brotman MA, Leibenluft E, McMahon FJ. Neurocognitive functioning in euthymic patients with bipolar disorder and unaffected relatives: A review of the literature. Neuroscience & Biobehavioral Reviews. 2016;69:193–215. doi:10.1016/j.neubiorev.2016.08.002.

27. Bora E. Neurocognitive features in clinical subgroups of bipolar disorder: A meta-analysis. Journal of Affective Disorders. 2018;229:125–134. doi:10.1016/j.jad.2017.12.057.

28. Keramatian K, Torres IJ, Yatham LN. Neurocognitive functioning in bipolar disorder: What we know and what we don’t. Dialogues in Clinical Neuroscience. 2021;23(1):29–38. doi:10.1080/19585969.2022.2042164.

29. Nitzburg GC, Cuesta-Diaz A, Ospina LH, Russo M, Shanahan M, Perez-Rodriguez M, et al. Organizational Learning Strategies and Verbal Memory Deficits in Bipolar Disorder. Journal of the International Neuropsychological Society : JINS. 2017;23(4):358–366. doi:10.1017/S1355617717000133.

30. Mundo E, Tharmalingham S, Neves-Pereira M, Dalton EJ, Macciardi F, Parikh SV, et al. Evidence that the N-methyl-D-aspartate subunit 1 receptor gene (GRIN1) confers susceptibility to bipolar disorder. Molecular Psychiatry. 2003;8(2):241–245. doi:10.1038/sj.mp.4001218.

31. Bundo M, Ueda J, Nakachi Y, Kasai K, Kato T, Iwamoto K. Decreased DNA methylation at promoters and gene-specific neuronal hypermethylation in the prefrontal cortex of patients with bipolar disorder. Molecular Psychiatry. 2021;26(7):3407–3418. doi:10.1038/s41380-021-01079-0.

32. Georgi A, Jamra RA, Schumacher J, Becker T, Schmael C, Deschner M, et al. No association between genetic variants at the GRIN1 gene and bipolar disorder in a German sample. Psychiatric Genetics. 2006;16(5):183. doi:10.1097/01.ypg.0000242194.36150.2b.

33. Catts VS, Lai YL, Weickert CS, Weickert TW, Catts SV. A quantitative review of the postmortem evidence for decreased cortical N-methyl-d-aspartate receptor expression levels in schizophrenia: How can we link molecular abnormalities to mismatch negativity deficits? Biological Psychology. 2016;116:57–67. doi:10.1016/j.biopsycho.2015.10.013.

34. McHugh TJ, Tonegawa S. CA3 NMDA receptors are required for the rapid formation of a salient contextual representation. Hippocampus. 2009;19(12):1153–1158. doi:10.1002/hipo.20684.

35. Ernst C, Deleva V, Deng X, Sequeira A, Pomarenski A, Klempan T, et al. Alternative Splicing, Methylation State, and Expression Profile of Tropomyosin-Related Kinase B in the Frontal Cortex of Suicide Completers. Archives of General Psychiatry. 2009;66(1):22–32. doi:10.1001/archpsyc.66.1.22.

36. Li N, He X, Zhang Y, Qi X, Li H, Zhu X, et al. Brain-derived neurotrophic factor signalling mediates antidepressant effects of lamotrigine. International Journal of Neuropsychopharmacology. 2011;14(8):1091–1098. doi:10.1017/S1461145710001082.

37. Hashimoto K. Brain-derived neurotrophic factor-TrkB signaling and the mechanism of antidepressant activity by ketamine in mood disorders. European Archives of Psychiatry and Clinical Neuroscience. 2020;270(2):137–138. doi:10.1007/s00406-020-01095-1.

38. Gideons ES, Lin PY, Mahgoub M, Kavalali ET, Monteggia LM. Chronic lithium treatment elicits its antimanic effects via BDNF-TrkB dependent synaptic downscaling. eLife. 2017;6:e25480. doi:10.7554/eLife.25480.

39. Wang Z, Fan J, Gao K, Li Z, Yi Z, Wang L, et al. Neurotrophic Tyrosine Kinase Receptor Type 2 (NTRK2) Gene Associated with Treatment Response to Mood Stabilizers in Patients with Bipolar I Disorder. Journal of Molecular Neuroscience. 2013;50(2):305–310. doi:10.1007/s12031-013-9956-0.

40. Bennett MR, Lagopoulos J. Stress and trauma: BDNF control of dendritic-spine formation and regression. Progress in Neurobiology. 2014;112:80–99. doi:10.1016/j.pneurobio.2013.10.005.

41. Szymanski J, Minichiello L. NKCC1 Deficiency in Forming Hippocampal Circuits Triggers Neurodevelopmental Disorder: Role of BDNF-TrkB Signalling. Brain Sciences. 2022;12(4):502. doi:10.3390/brainsci12040502.

42. Haukvik UK, Gurholt TP, Nerland S, Elvsåshagen T, Akudjedu TN, Alda M, et al. In vivo hippocampal subfield volumes in bipolar disorder—A mega-analysis from The Enhancing Neuro Imaging Genetics through Meta-Analysis Bipolar Disorder Working Group. Human Brain Mapping. 2022;43(1):385–398. doi:10.1002/hbm.25249.

43. Chadwick MJ, Bonnici HM, Maguire EA. CA3 size predicts the precision of memory recall. Proceedings of the National Academy of Sciences. 2014;111(29):10720–10725. doi:10.1073/pnas.1319641111.

44. Zhang ZJ, Reynolds GP. A selective decrease in the relative density of parvalbumin-immunoreactive neurons in the hippocampus in schizophrenia. Schizophrenia Research. 2002;55(1):1–10. doi:10.1016/S0920-9964(01)00188-8.

45. Konradi C, Zimmerman EI, Yang CK, Lohmann KM, Gresch P, Pantazopoulos H, et al. Hippocampal Interneurons in Bipolar Disorder. Archives of General Psychiatry. 2010;68(4):340. doi:10.1001/archgenpsychiatry.2010.175.

46. Mertens J, Wang Q, Kim Y, Yu D, Pham S, Yang B, et al. Differential responses to lithium in hyperexcitable neurons from patients with bipolar disorder. Nature. 2015;527(7576):95–99. doi:10.1038/nature15526.

47. Stern S, Sarkar A, Stern T, Mei A, Mendes APD, Stern Y, et al. Mechanisms Underlying the Hyperexcitability of CA3 and Dentate Gyrus Hippocampal Neurons Derived From Patients With Bipolar Disorder. Biological Psychiatry. 2020;88(2):139–149. doi:10.1016/j.biopsych.2019.09.018.

48. Stern S, Sarkar A, Galor D, Stern T, Mei A, Stern Y, et al. A Physiological Instability Displayed in Hippocampal Neurons Derived From Lithium-Nonresponsive Bipolar Disorder Patients. Biological Psychiatry. 2020;88(2):150–158. doi:10.1016/j.biopsych.2020.01.020.

49. Rebola N, Carta M, Mulle C. Operation and plasticity of hippocampal CA3 circuits: implications for memory encoding. Nature Reviews Neuroscience. 2017;18(4):208–220. doi:10.1038/nrn.2017.10.

50. Santos R, Linker SB, Stern S, Mendes APD, Shokhirev MN, Erikson G, et al. Deficient LEF1 expression is associated with lithium resistance and hyperexcitability in neurons derived from bipolar disorder patients. Molecular Psychiatry. 2021;doi:10.1038/s41380-020-00981-3.

51. Stern S, Santos R, Marchetto M, Mendes A, Rouleau G, Biesmans S, et al. Neurons derived from patients with bipolar disorder divide into intrinsically different sub-populations of neurons, predicting the patients’ responsiveness to lithium. Molecular Psychiatry. 2018;23(6):1453–1465. doi:10.1038/mp.2016.260.

52. Tripathi U, Mizrahi L, Alda M, Falkovich G, Stern S. Information theory characteristics improve the prediction of lithium response in bipolar disorder patients using a support vector machine classifier. Bipolar Disorders. 2023;25(2):110–127. doi:10.1111/bdi.13282.

